# Lipid droplet dynamics during the budding yeast cell cycle influence the timing of cell cycle START

**DOI:** 10.1101/2025.09.25.678471

**Authors:** Hanna M. Terpstra, Katerina Loukogiannaki, Vasiliki Tsousi, Matthias Heinemann

## Abstract

Recent work has revealed that metabolism is dynamic over the budding yeast cell cycle, showing that the NAD(P)H autofluorescence oscillates, and protein and lipid biosynthesis are dynamic. Dynamic storage and liquidation of neutral lipids over the cell cycle could contribute to these metabolic dynamics. However, the dynamics of neutral lipids over the cell cycle are as yet unknown. To elucidate them, we established mNeonGreen-Tgl3 and Pln1-mNeonGreen as protein markers for lipid droplets (LDs) and determined LD dynamics during the cell cycle with single-cell time-lapse microscopy. We found oscillations in the LD number over the cell cycle, with a notable trough around START. Deletion of the genes responsible for either the synthesis of triacylglycerol (TAG) or its mobilisation from LDs lowered LD numbers and abolished the oscillation in LD number. Moreover, in these deletion mutants, we found START to be delayed, suggesting that the mobilisation of TAG from LDs is required for its timely occurrence. The influence of LD dynamics on the timing of START emphasises that research studying cell cycle commitment should consider storage lipid metabolism as a potential contributor to cell cycle START.

## INTRODUCTION

Proper control of cell growth and division is fundamental for cells. In eukaryotes, the cyclin/cyclin-dependent kinase (CDK) machinery is considered the primary regulator of this process (Mendenhall & Hodge, 1998). However, recent work in *S. cerevisiae* suggests that an intrinsically dynamic metabolism could also play a role in cell cycle regulation. For instance, CDK-independent oscillations in NAD(P)H levels were found essential for cell division (Papagiannakis et al., 2017) and an increase in protein biosynthesis during the G1 phase was found to drive commitment to the cell cycle by temporarily increasing Cln3 levels (Litsios et al., 2019). As the dynamics of metabolism evidently are also involved in cell cycle control, it is important to precisely understand the metabolic dynamics and their underlying cause in order to investigate the precise mechanisms through which metabolism controls cell cycle progression.

Multiple metabolic processes oscillate along the cell cycle: the glycolytic flux peaks between cytokinesis and late G1 phase (Monteiro et al., 2019) and mobilisation of stored carbohydrates occurs during G1 (Müller et al., 2003; Silljé et al., 1997; Zhao et al., 2016). Lipid biosynthesis peaks during S/G2/M (Takhaveev et al., 2023), coinciding with increased protein levels of the biosynthetic enzymes for both ergosterol (Blank et al., 2020) and fatty acids (Blank et al., 2017) and increased triacylglycerol levels (Blank et al., 2020) in this phase.

Besides lipid synthesis, lipid storage could also be dynamic during the cell cycle. Neutral lipids, namely triacylglycerols (TAG) and steryl esters (SE), are stored in lipid droplets (LDs), which are critical during growth transitions. When budding yeast enters stationary phase, TAGs are synthesised and stored in LDs (Czabany et al., 2008; Markgraf et al., 2014). In contrast, when yeast cells exit stationary phase, these neutral lipids are mobilised and LD size and number decrease (Ganesan et al., 2020; Markgraf et al., 2014; Ouahoud et al., 2018). Furthermore, mutants deficient in TAG mobilisation resume growth more slowly than the wild type when cells grown to stationary phase are shifted into fresh medium (Kurat et al., 2009; Ouahoud et al., 2018). Based on these findings, we wondered whether neutral lipid storage could also be dynamic during the cell cycle and if so, whether these dynamics would affect cell cycle progression.

In this study, we established protein markers for LDs and used them to uncover cell cycle oscillations of the number of LDs. These oscillations show a trough around cell cycle START and a peak midway through S/G2/M, indicating that LDs are first depleted and then re-formed as the cell cycle progresses. In strains lacking enzymes for TAG synthesis or mobilisation, the dynamics of the LD number are lost and START is delayed. Our findings indicate that mobilisation of TAG from LDs contributes to the timely occurrence of START, suggesting that LD dynamics play a role in cell cycle control.

## RESULTS

### Establishing protein markers for dynamic, dye-free identification of lipid droplets

To elucidate the dynamic behaviour of lipid droplets during the cell cycle in *S. cerevisiae*, we used fluorescence microscopy in combination with a microfluidic setup (Huberts et al., 2013; Lee et al., 2012), which allows the generation of dynamic single-cell data over several cell cycles under constant growth conditions. To visualise LDs, we did not use chemical stains, since this would have required the continuous addition of a dye to the growth medium to counteract the dilution of the dye as cells grow. Instead, to achieve dye-free identification of LDs, we chose to tag proteins of the LD proteome (Grillitsch et al., 2011) that localise to LDs and no other organelles (SGD, 2017), with mNeonGreen and use these tagged proteins as reporters for LDs. Using image databases of *S. cerevisiae* with GFP-tagged proteins (Huh et al., 2003; Weill et al., 2018), we selected proteins of the LD proteome whose LD localisation remained unchanged after introduction of a fluorescent tag. Specifically, we selected Pln1, Tgl3 and Rrt8 as candidate protein markers for LDs. Pln1 stimulates the accumulation of TAG and stabilises LDs (Gao et al., 2017). Tgl3 is a TAG lipase and thus mobilises neutral lipids from LDs (Athenstaedt & Daum, 2003; Chauhan et al., 2015) and Rrt8 has been implicated in spore wall assembly (Lin et al., 2013) and transport of plasma membrane proteins (Ueno et al., 2016).

To verify that these candidate reporter proteins indeed localise to LDs and can thus serve as LD markers, we fixed cells with formaldehyde, stained LDs with the red fluorescent dye BODIPY-TR and then assessed colocalisation between the stained LDs and the mNeonGreen-tagged marker proteins. Given that the chosen marker proteins have different LD-related functions, we considered that certain marker proteins might localise only to a subset of all LDs, or reversely, could have a localised function beyond LDs.

Before testing the colocalisation between LDs detected with BODIPY-TR and the marker proteins, we first ensured optimal alignment between images recorded in the green and red fluorescence channels using a bilinear transformation approach. As LDs have an average diameter of 0.4 µm (Czabany et al., 2008), corresponding to less than three pixels in our microscopy setup, even minor misalignment between the two fluorescence channels would have significantly distorted the colocalisation analysis.

We then used wild-type cells stained with BODIPY-TR to show that the red BODIPY-TR signal is not detected in the GFP channel **(Figure S1A-D)**. Furthermore, we compared the fluorescence intensity measured in the GFP channel between a wild-type control, *i.e.* autofluorescence, and the three LD reporter strains and verified that bright spots of autofluorescence occasionally present in GFP images of wild-type cells are not detectable in the LD reporter strains with the applied spot detection algorithm (Terpstra et al., 2024) and detection settings **(Figure S1E-G)**. The following colocalisation analysis therefore is not confounded by the detection of GFP puncta that are due to autofluorescence instead of an LD marker protein, or artefacts of BODIPY-TR fluorescence detected in the GFP channel.

Next, to perform the actual colocalisation analysis, we imaged cells stained with BODIPY-TR that also expressed one of the LD reporter fusion proteins Pln1-mNG, mNG-Tgl3 and Rrt8-mNG. We detected puncta of the mNeonGreen-tagged LD marker proteins in the GFP channel and LDs stained with BODIPY-TR in the RFP channel and saw that puncta of the reporter proteins and puncta identified with BODIPY-TR were often found at similar locations within the cell **(Figure 1A)**. To quantify colocalisation between puncta of BODIPY-TR and puncta of the LD marker protein candidates, we differentiated between ‘certain’ and ‘ambiguous’ colocalisation. Colocalisation was classified as ‘certain’ if puncta in the two imaging channels had identical midpoints or were located at most one pixel apart; ‘ambiguous’ colocalisation refers to puncta whose midpoints were at most two pixels apart. In the following, we consider both the ‘certain’ and ‘ambiguous’ categories as colocalised.

**Figure 1.**
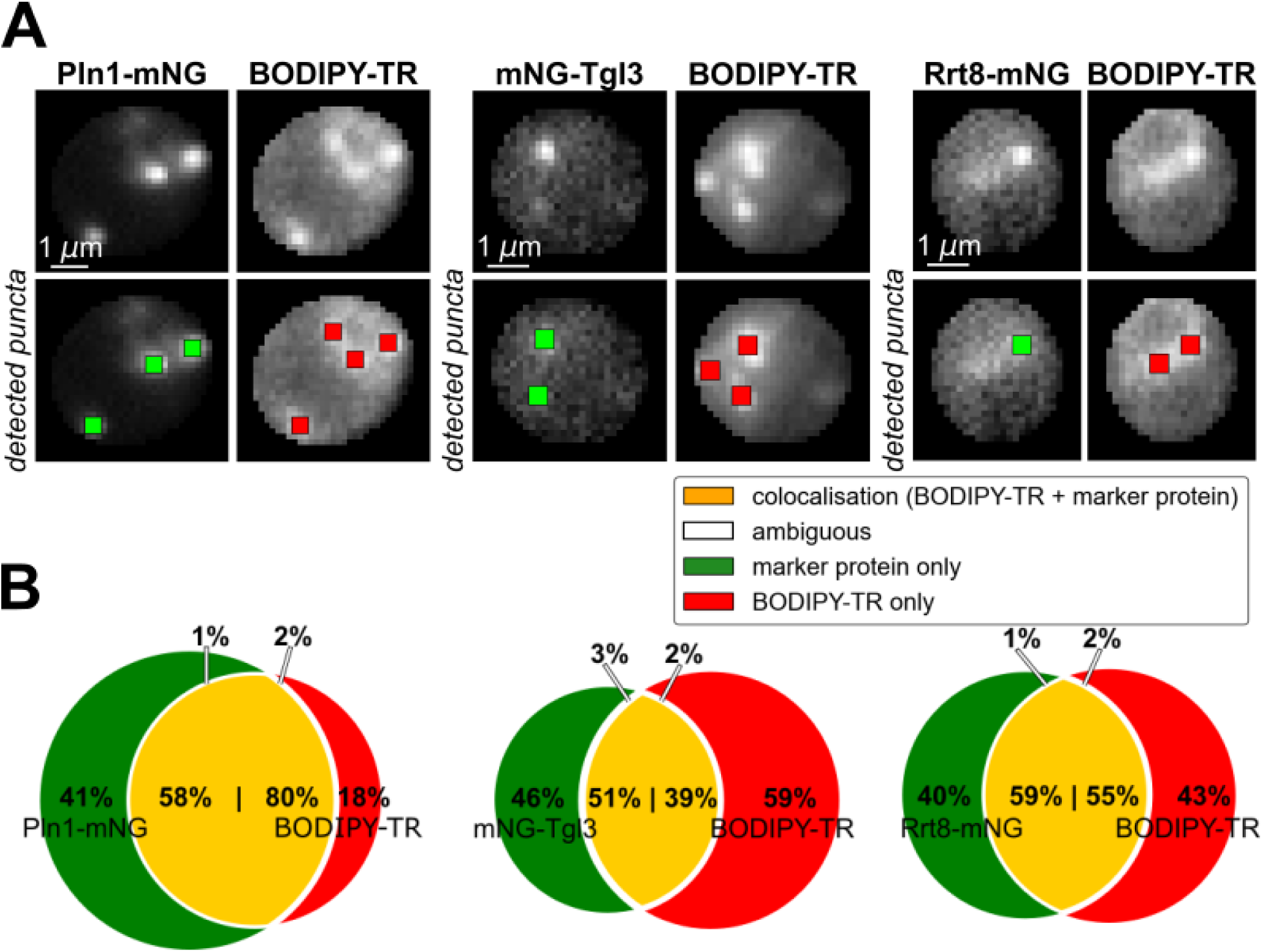
Establishing protein markers for dynamic, dye-free identification of lipid droplets. **(A)** Cells expressing a candidate LD reporter protein tagged with mNeonGreen (Pln1-mNG, mNG-Tgl3 or Rrt8-mNG) were fixed with formaldehyde, stained with BODIPY-TR to visualise LDs and then imaged (upper row). Puncta of marker proteins and LDs stained with BODIPY-TR were detected with a spot detection algorithm (lower row); **(B)** Colocalisation between LDs stained with BODIPY-TR and puncta of the three candidate LD reporter proteins Pln1-mNG, mNG-Tgl3 and Rrt8-mNG. The left circle of each Venn diagram represents the marker protein puncta identified in the GFP channel image and the right circle represents puncta of BODIPY-TR in the RFP channel image. The orange overlapping region between the two circles represents the colocalised puncta. The white regions represent puncta whose colocalisation is ambiguous: while puncta in the two fluorescence channels have midpoints close together, their colocalisation is likely, but not certain. The following number of cells and puncta were assessed in the colocalisation analysis: Pln1-mNG: 153 cells, 304 BODIPY-TR puncta, 424 reporter protein puncta; mNG-Tgl3: 217 cells, 345 BODIPY-TR puncta, 263 reporter protein puncta; Rrt8-mNG: 245 cells, 323 BODIPY-TR puncta, 302 reporter protein puncta.

We first assessed the fraction of BODIPY-TR puncta that colocalise with a punctum of an LD reporter protein. We found that 82%, 57% and 41% of BODIPY-TR puncta have a corresponding punctum of Pln1-mNG, Rrt8-mNG and mNG-Tgl3, respectively **(Figure 1B)**. We also determined the fraction of reporter protein puncta that colocalise with a BODIPY-TR punctum and found that 60%, 59% and 54% of protein puncta colocalise with a punctum of a stained LD for the marker proteins Rrt8-mNG, Pln1-mNG and mNG-Tgl3 **(Figure 1B)**. These data show that not all detected LDs stained with BODIPY-TR are also visible as a marker protein punctum, consistent with the existence of subpopulations of LDs with distinct proteomes (Eisenberg-Bord et al., 2018) and lipid content (Meyers et al., 2016). Furthermore, the data show that not all puncta of the LD reporter proteins colocalise with an LD stained with BODIPY-TR. Protein puncta of Pln1 without a corresponding BODIPY-TR puncta could be LDs that are still too small for detection with BODIPY-TR, consistent with the finding that appearance of Pln1 puncta often precedes BODIPY-TR puncta by a few minutes (Gao et al., 2017). The Rrt8-mNG and mNG-Tgl3 puncta without a corresponding BODIPY-TR punctum might be indicative of alternative functions of these proteins, outside LDs.

Overall, our results show that LDs are heterogeneous, with different marker proteins likely representing distinct subsets of LDs. It is conceivable that one reporter could miss a subset of LDs that another reporter could identify. As all three tested LD markers are endogenous proteins with their own biological functions, it is also possible that their dynamics in part reflect other biological functions, besides reporting the dynamics of LDs. Finally, as the localisation of Pln1-mNG is most comparable to that of the LD dye BODIPY-TR, Pln1-mNG is the most suitable LD marker to investigate whether there are LD dynamics during the cell cycle.

### Number of lipid droplets oscillates over the cell cycle

Next, we performed time-lapse microscopy experiments to determine the number and size of LDs over the cell cycle using Pln1-mNG and mNG-Tgl3 as markers for LDs. Notably, we did not employ Rrt8-mNG as an LD reporter in our time-lapse experiments. Due to the lower expression of Rrt8 compared to Pln1 and Tgl3 (Breker et al., 2013; Chong et al., 2015; SGD, 2017), imaging required higher light exposure, resulting in prolonged cell cycle durations **(Figure S2)**, likely due to phototoxicity. By studying LD dynamics with two distinct protein reporters, we aimed to elucidate the more general cell cycle dynamics of LDs, independent of specific functions of the reporter proteins. Therefore, we focused on global similarities in any eventual dynamics observed with the two markers, disregarding small differences between the two reporters.

We used an automated spot detector algorithm (Terpstra et al., 2024) to detect LDs and to estimate their size in the time-lapse images. We normalised the number of LDs detected within a cell to the area of its cross-section. To project the detected LD characteristics on a common cell cycle progression coordinate, we aligned all LD cell cycle trajectories for mitotic exit, START and budding. Finally, we applied Gaussian process regression to predict the average cell cycle dynamics of the LD number normalised to the cell cross-area, and of the LD size.

With both LD reporter proteins, we found that the area-normalised LD number oscillates over the cell cycle with a minimum around START and a maximum between budding and mitotic exit **(Figure 2A-B)**. Density plots showing all data points reveal the same cell cycle dynamics **(Figure S3A-B)** as the population average dynamics predicted with Gaussian process regression. With Pln1-mNG a second, less pronounced trough is visible late in the second half of the cell cycle **(Figure 2A)**. In contrast to the area-normalised LD number, LD size is constant throughout the cell cycle **(Figure 2C-D)**. Again, density plots confirm the dynamics predicted with Gaussian process regression **(Figure S3C-D)**. The cell cycle dynamics of the summed sizes of all LDs per cell, which notably is not normalised to the cell cross-area, strongly resemble the dynamics of the area-normalised LD number **(Figure S3E-F)**. The similarity between these dynamics indicates that the oscillation of the LD number is not an artefact resulting from the normalisation to the cell cross-area.

**Figure 2.**
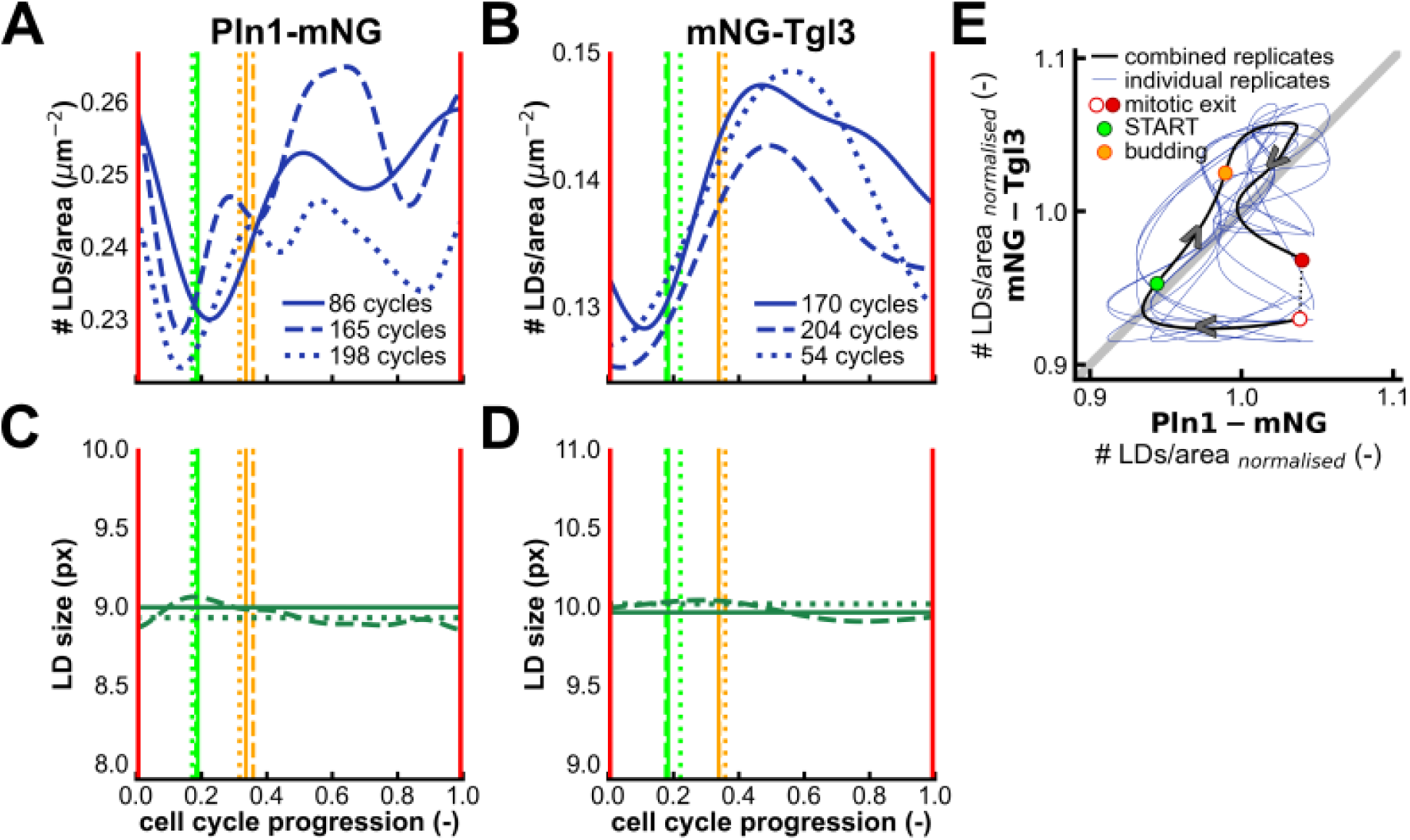
Number of lipid droplets oscillates over the cell cycle while LD size is constant. LDs were identified in time-lapse microscopy images of cells expressing either Pln1-mNG (A, C) or mNG-Tgl3 (B, D) as an LD marker. For each LD reporter strain, three biological replicates, whose results are represented by different line styles, were performed. Cell cycle trajectories were aligned from one mitotic exit to the next (red vertical lines at cell cycle progression values 0 and 1) and for occurrence of START (bright green vertical line) and budding (orange vertical line) between these mitotic exit events. Gaussian process regression was used to predict population averages of **(A-B)** area-normalised number of detected LDs and **(C-D)** size of the detected LDs; **(E)** To compare the dynamics of area-normalised number of LDs as observed with the reporter proteins Pln1-mNG and mNG-Tgl3, we normalised every Gaussian process regression result to its own average value and plotted results obtained with mNG-Tgl3 against those obtained with Pln1-mNG. The grey line indicates y = x. Thin coloured lines denote the combination of individual replicates using the two LD reporters (3x3 combinations). The thick black line indicates combined results of the three replicate experiments performed with each LD reporter and was obtained by averaging the three Gaussian process regression outputs for every timepoint. The circular markers on the black curve denote the occurrence of mitotic exit, START and budding.

We also noticed that the area-normalised LD number determined with mNG-Tgl3 is almost twofold lower than that measured with Pln1-mNG **(Figure 2A-B)**. This difference could be explained by the localisation of mNG-Tgl3 to a specific subset of LDs. First, when LD formation in a mutant strain is induced by expression of the diacylglycerol acyltransferase Dga1, Tgl3 is not detected on newly formed LDs for the first hour after induction (Gao et al., 2017). Second, since Tgl3 is an enzyme that uses TAG as its substrate (Athenstaedt & Daum, 2003; Rajakumari & Daum, 2010) and subpopulations of TAG-specific LDs have been reported in mammalian cells (Hsieh et al., 2012) and *S. pombe* (Meyers et al., 2016), it is plausible that Tgl3 would localise only to TAG-enriched LDs, while Pln1 does not show this specific localisation.

To investigate the similarities and differences of the oscillations observed with the two LD reporters, we averaged the three replicates performed with each reporter, normalised the result to its mean and plotted the normalised trajectories obtained with Pln1-mNG and mNG-Tgl3 against each other. Here, we found a positive correlation from shortly before START until the final 15% of the cell cycle **(Figure 2E)**, indicating that the two markers report comparable dynamics of the area-normalised LD number during this time interval. In contrast, the dynamics are different around mitotic exit. This difference could arise from the different functions of the two LD reporter proteins. Specifically, since Pln1 is important for the stabilisation of nascent LDs (Gao et al., 2017), the increase in area-normalised LD number observed with Pln1-mNG at the end of the cell cycle could indicate the formation of new LDs. An increase in LD numbers at the end of the cell cycle would be consistent with the above-average lipid biosynthetic activity at the end of the cell cycle after its peak in the middle of S/G2/M (Takhaveev et al., 2023) and the storage of triacylglycerol during septation (P. L. Yang et al., 2016). If neutral lipids are mobilised from these same LDs after mitotic exit, leading to their disappearance, this would explain the more pronounced drop in area-normalised LD number observed with Pln1-mNG compared to mNG-Tgl3.

Overall, employing two distinct LD reporter proteins, we have elucidated the general cell cycle dynamics of LDs, which suggest that neutral lipids are mobilised from LDs between mitotic exit and START, as evidenced by a trough in the area-normalised LD number, while the neutral lipid stores are replenished during S/G2/M. These findings demonstrate that LDs are not stagnant organelles containing energy reserves to be used in case of nutrient shortages. Instead, their content is mobilised and re-synthesised as cells go through the cell cycle.

### Dynamic LD fluorescence intensity likely results from LD number dynamics

After we discovered the cell cycle dynamics of the LD number, we asked whether the fluorescence intensity measured on those LDs was also dynamic. Studying the dynamics of the fluorescence intensity of the LD reporters Pln1-mNG and mNG-Tgl3 on LDs and in the cytoplasm could help us distinguish between observations that reflect the general dynamics of LDs along the cell cycle and observations that relate to the distinct biological functions of the LD marker proteins. We assessed the cell cycle dynamics of the fluorescence intensity measured inside the whole cell mask, on LDs and in the cytoplasm. The latter was determined by excluding all pixels belonging to an LD and calculating the average intensity of all other pixels within the cell mask. While the fluorescence intensity inside the whole cell and in the cytoplasm was found to be almost constant, the fluorescence intensity on LDs oscillates along the cell cycle **(Figure 3A-B)**. With both Pln1-mNG and mNG-Tgl3, the fluorescence intensity on LDs peaked around START and was minimal during S/G2/M.

**Figure 3.**
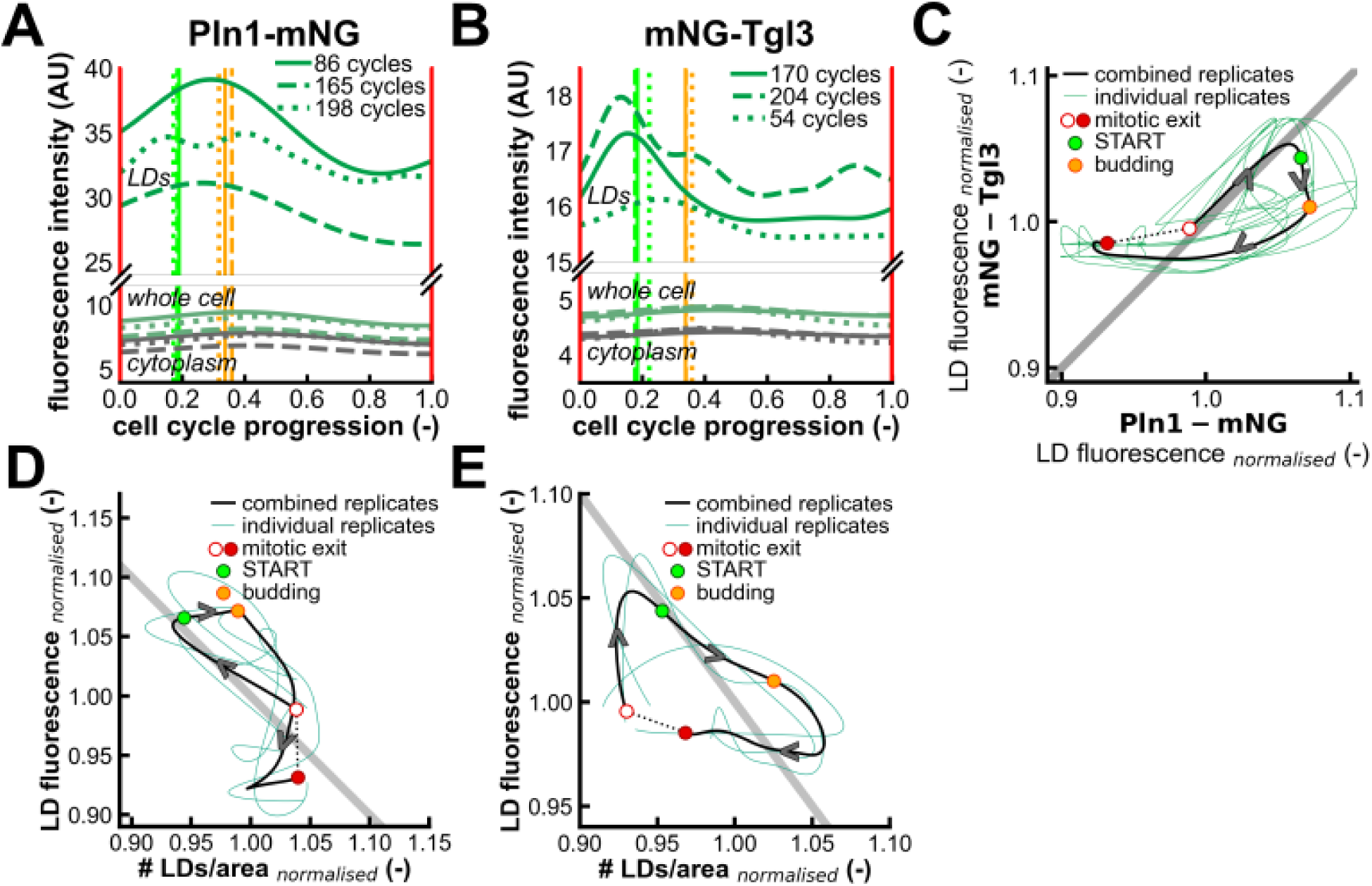
Dynamic LD fluorescence intensity anticorrelates with area-normalised LD number. LDs were identified in time-lapse microscopy images of cells expressing either Pln1-mNG (A, D) or mNG-Tgl3 (B, E) as an LD reporter protein. For each reporter strain, multiple biological replicates, whose results are represented by different line styles, were performed. Cell cycle trajectories were aligned from one mitotic exit to the next (red vertical lines at cell cycle progression values 0 and 1) and for the occurrence of START (bright green vertical line) and budding (orange vertical line) between the two subsequent instances of mitotic exit. Gaussian process regression was used to predict population averages of **(A-B)** average fluorescence intensity of detected LDs, the cytoplasmic region and the whole cell mask; **(C)** To allow direct comparison between the cell cycle dynamics of the fluorescence intensity of LDs detected with Pln1-mNG or mNG-Tgl3, we normalised every Gaussian process regression output to its own mean value and plotted the trajectories obtained with mNG-Tgl3 against those obtained with Pln1-mNG. The grey line indicates y = x, on which all data points would lie in case of perfect correlation. Thin coloured lines denote combinations of individual replicates (3x3 combinations). The thick black line represents the averaged results of the three replicates performed with each LD reporter protein. The circular markers on this curve represent the occurrence of the cell cycle events mitotic exit, START and budding; **(D-E)** To show anticorrelation between the area-normalised LD number and the fluorescence intensity measured on LDs, we normalised every Gaussian process output to its own mean and plotted the results against each other. Thin coloured lines denote the results for the individual replicates and the thick black line represent averaged results of the three replicates; circular markers on this black trajectory denote mitotic exit, START and budding. The grey line indicates where data points would lie in case of perfect anticorrelation.

To compare the dynamics of LD fluorescence measured with the two reporters and to distinguish between general and reporter protein-specific behaviour, we normalised the three replicates performed with each marker protein to their respective means, took the average of the three replicates and plotted the resulting trajectory for mNG-Tgl3 against that for Pln1-mNG. We found that the trajectories obtained with the two LD marker proteins correlate positively **(Figure 3C)**, demonstrating that, along most of the cell cycle, the oscillations of the fluorescence intensities measured with the two reporters are comparable. However, this correlation is absent during the last stretch of the cell cycle, when the fluorescence of mNG-Tgl3 on LDs is almost constant, while the fluorescence of Pln1-mNG on LDs is still decreasing. The sharp peak of mNG-Tgl3 intensity on LDs around START could be related to its function as a triacylglycerol lipase, as the decrease in the area-normalised LD number between mitotic exit and START **(Figure 2A-B)** indeed indicates that neutral lipids are mobilised from LDs before START.

Comparing the dynamics of the area-normalised LD number **(Figure 2A-B)** and the LD fluorescence intensity **(Figure 3A-B)**, we noticed that the minimum of the area-normalised LD number around START coincides with the maximum of the LD fluorescence intensity while, vice versa, the maximum of the area-normalised LD number and the minimum in the fluorescence intensity on LDs both occur during S/G2/M. Indeed, when we plotted the area-normalised LD number against the LD fluorescence, each cell cycle trajectory normalised to its own mean, we saw an anticorrelation between these two characteristics. This anticorrelation was observed both with Pln1-mNG and with mNG-Tgl3 as an LD reporter protein **(Figure 3D-E)**.

To comprehend why the fluorescence intensity of LDs would anticorrelate with the LD number, we aimed to unite all observed LD characteristics, *i.e.* number, size and fluorescence intensity as well as the concentration of the fluorescent reporter protein, in a unified explanation. First, we excluded that a dynamic concentration of LD marker protein causes the cell cycle dynamics of the LD fluorescence intensity. The concentration, proxied by the average fluorescence intensity within the cell mask, is almost constant for both Pln1-mNG and mNG-Tgl3 **(Figure 3A-B)** and thus does not elicit the dynamic LD fluorescence intensity. Second, changes in partitioning of the LD reporter proteins between LDs and the cytoplasm cannot drive the changes in LD fluorescence either. If this partitioning changed, fluorescence on LDs and in the cytoplasmic would show opposite trends, since marker protein moving from cytoplasm to LDs would cause the cytoplasmic fluorescence intensity to decrease and the fluorescence intensity of LDs to increase, and vice versa. However, the fluorescence intensity on LDs is dynamic, while the fluorescence in the cytoplasm is constant **(Figure 3A-B)**. Third, we considered that LD size could account for changes in the fluorescence intensity of LDs. If the amount of marker protein localised to an LD stayed constant while that LD increased in size, the marker protein would be dispersed and the fluorescence intensity would decrease. Conversely, if the LD shrunk, the marker protein would become more concentrated, increasing the fluorescence intensity. However, LD size is constant **(Figure 2C-D)**, so changes in LD size do not cause the dynamic fluorescence intensity of LDs. Lastly, the dynamic area-normalised LD number **(Figure 2A-B)** and its anticorrelation with the fluorescence intensity of LDs **(Figure 3D-E)** can drive the changes in the LD fluorescence intensity. When the number of LDs is low, the total pool of reporter protein is spread across few LDs, resulting in high fluorescence intensities on those LDs. When the number of LDs increases, the same reporter protein molecules are distributed over more LDs, and consequently, the fluorescence intensity on the LDs will be lower. Thus, the observed changes in LD number could be responsible for the dynamics in LD fluorescence intensity, whereby the anticorrelation between fluorescence intensity of LDs and the area-normalised LD number would be explained.

Overall, our results show that the fluorescence intensity of LDs oscillates along the cell cycle and anticorrelates with the area-normalised number of LDs. In contrast, reporter protein concentrations, the partitioning of reporter proteins between LDs and the cytoplasm, and LD size are constant. Together, these findings indicate that the cell cycle dynamics of the LD fluorescence intensity are likely driven by the oscillating LD number.

### TAG storage and mobilisation give rise to LD dynamics

After we discovered the cell cycle dynamics of LD number and fluorescence intensity, we wondered whether we could identify which biological process is responsible for the oscillations. Changes in the LD number and fluorescence could be due to changing synthesis and mobilisation of TAG and steryl esters, as well as fission and fusion of LDs. To test whether TAG metabolism was responsible for the LD cell cycle dynamics, we perturbed TAG metabolism and subsequently observed the area-normalised LD number along the cell cycle. To perturb TAG synthesis, we deleted the genes encoding the two major TAG synthases in *S. cerevisiae* Lro1 (Oelkers et al., 2000) and Dga1 (Oelkers et al., 2002). In a separate strain, we deleted the genes encoding the TAG lipases Tgl3 (Athenstaedt & Daum, 2003) and Tgl4 (Athenstaedt & Daum, 2005), thereby blocking the mobilisation of TAG from LDs. Notably, in the cells with perturbed TAG metabolism, we only used Pln1-mNG as an LD reporter, since our other reporter protein, Tgl3, was deleted in one of the TAG mutant strains.

Before investigating the LD cell cycle dynamics in the two double deletion strains, we first assessed how the perturbation of TAG metabolism affected the LD phenotype independent of the cell cycle stage. To this end, we analysed fluorescence microscopy snapshots of cells from exponential cultures. We saw that fluorescence intensity of Pln1-mNG both within the entire cell mask and on LDs was higher in *ΔTGL4ΔTGL3* compared to the wild type, and lower in *ΔDGA1ΔLRO1* **(Figure S4)**. These changes in the expression of Pln1, which is important for the formation and stabilisation of LDs, imply that the number and size of LDs could also be different in the two mutation strains. Indeed, we found that the area-normalised LD number was significantly lower in both deletion strains relative to the wild type **(Figure 4A)**, with no puncta detected in 44% of Δ*DGA1*Δ*LRO1* cells and 11% of Δ*TGL3*Δ*TGL4* cells. Moreover, the LDs in both deletion strains were bigger than those in the wild type **(Figure 4B)**. Therefore, perturbation of TAG metabolism, preventing its synthesis or mobilisation, affects both the number of LDs and their size.

**Figure 4.**
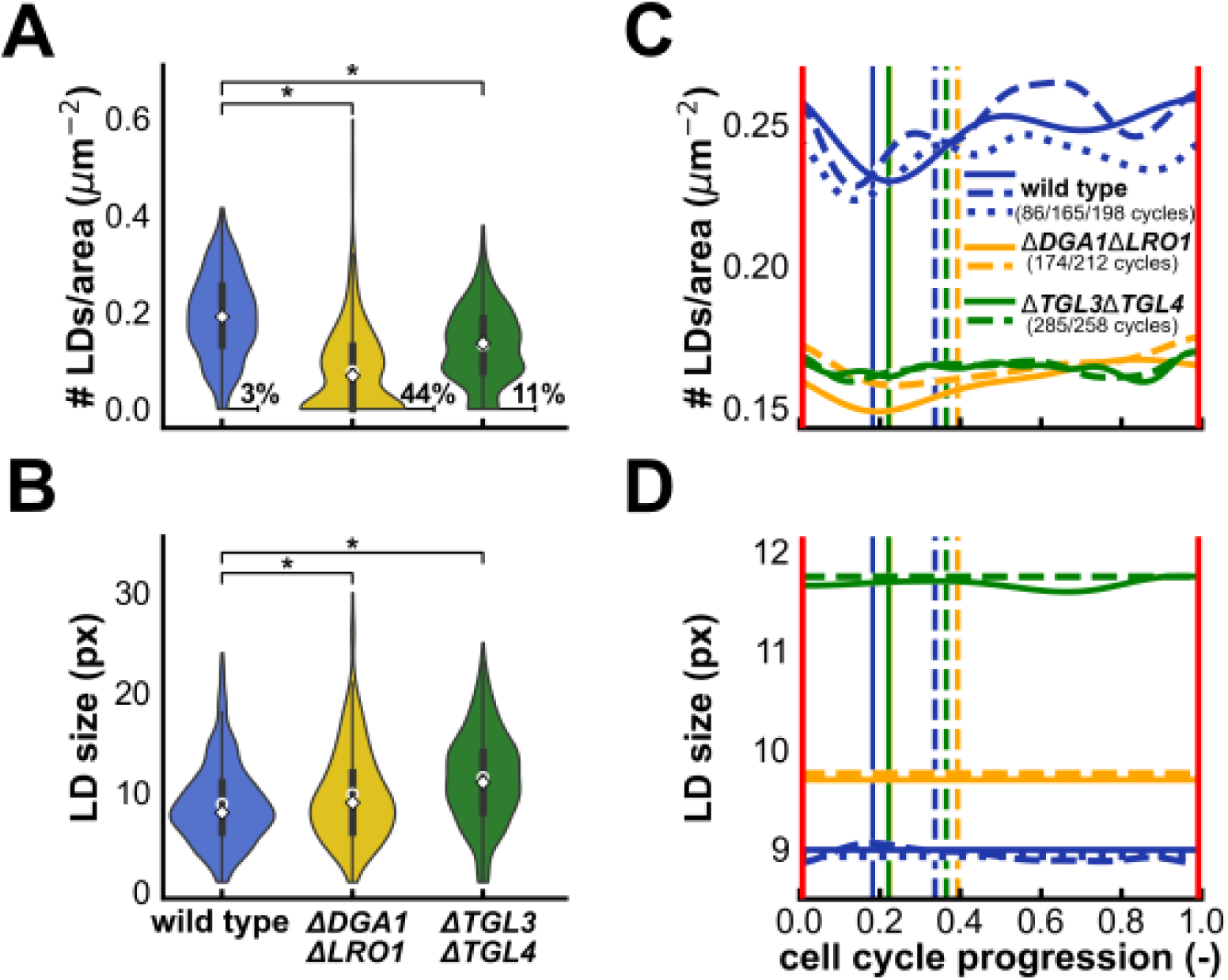
TAG storage and mobilisation give rise to LD dynamics. **(A-B)** Area-normalised number of LDs and LD size as determined from snapshots of cells from exponential cultures that express Pln1-mNG as an LD reporter in wild type, *ΔDGA1ΔLRO1* and *ΔTGL3ΔTGL4* (298, 314 and 264 cells, respectively). The median is indicated with a white diamond and an open circle indicates the mean. In both deletion backgrounds, the number of detected LDs per cell cross-area is significantly lower and LDs are significantly larger than in the wild type (two-sided Mann-Whitney U-test; p < 0.05). The percentage of cells without any detected LDs is indicated next to each violin in A; **(C-D)** Cell cycle dynamics of the area-normalised LD number and size of detected LDs predicted with Gaussian process regression applied to cell cycle-aligned single-cell trajectories from three biological backgrounds, indicated in different colours. Biological replicates are indicated by different line styles. Cell cycles were aligned from one occurrence of mitotic exit to the next (red vertical lines at cell cycle progression values 0 and 1) and for occurrence of START (solid vertical lines) and budding (dashed vertical lines).

Some of the changes in the LD phenotype may at first glance seem counterintuitive, but are in line with previous research. In the *ΔDGA1ΔLRO1* mutant, which is unable to synthesise TAG, we found larger LDs than in the wild type **(Figure 4B)** while one may expect a decrease in LD size due to the absence of TAG. The stabilisation of specifically small LDs by diacylglycerol acyltransferases (Kovacs et al., 2021; Wilfling et al., 2013) could explain the increased size of LDs in *ΔDGA1ΔLRO1*. The other deletion strain, *ΔTGL3ΔTGL4* can synthesise TAG, but cannot mobilise it from LDs. Therefore, one would not expect the observed decrease in LD number compared to the wild type **(Figure 4A)**. As Tgl4 is involved in the stabilisation of nascent LDs (Wang et al., 2024), its absence in *ΔTGL3ΔTGL4* may hinder LD formation, resulting in lower LD numbers compared to the wild type. Overall, these findings show that deleting the genes encoding the enzymes that synthetise or mobilise TAG changes LD morphology, and can have counterintuitive effects due to additional functions of these enzymes.

Next, we performed time-lapse microscopy experiments to investigate the cell cycle dynamics of LDs in the deletion strains and thereby determine whether TAG metabolism contributes to LD dynamics. In both deletion backgrounds, the cell cycle oscillations of the area-normalised LD number as observed in the wild type were lost **(Figure 4C)**. Similarly to the wild type, LD size was constant along the cell cycle in both deletion backgrounds **(Figure 4D)**. As LD fission would lead to the appearance of two smaller LDs originating from one larger LD, while LD fusion would have the opposite effect, the constant LD size along the cell cycle makes it improbable that LD fission and fusion contribute to the cell cycle dynamics of the LD number. In contrast, the constant area-normalised LD number in mutants unable to synthesise or mobilise TAG suggests that TAG metabolism must give rise to the LD dynamics as observed in wild-type cells. Hereby, we have established that TAG metabolism, instead of the storage and mobilisation of steryl esters or the fission and fusion of LDs, underlies the oscillations of the LD number during the cell cycle.

### Perturbing LDs through TAG metabolism delays START

Finally, we asked if the LD dynamics, as observed in the wild type but lost in the strains with perturbed TAG metabolism, could affect cell cycle progression. Since START is delayed when cells that cannot mobilise TAG from LDs resume growth after starvation (Kurat et al., 2009), we wondered if the same could be true in exponentially growing cells. To investigate this, we assessed the interrelation between the duration of the whole cell cycle and the time between mitotic exit and START in individual cell cycles. We plotted the duration of the mitotic exit to START phase against the whole cell cycle length in *ΔDGA1ΔLRO1*, *ΔTGL3ΔTGL4* and the wild type and performed linear regression to obtain equations that describe their interrelation **(Figure 5A)**. The regression lines describe the duration of mitotic exit to START as a function of cell cycle length and therefore, their slopes indicate the fraction of the cell cycle taken up by mitotic exit to START. We found that the slopes obtained from both deletion strains were steeper than those from the wild type **(Figure 5B)**, which means that mitotic exit to START takes up a larger fraction of the cell cycle in TAG mutants compared to the wild type. The change in the fraction of the cell cycle taken up by mitotic exit to START detected on the single-cell cycle level was not accompanied by extensive population-level changes in absolute duration of the cell cycle or its phases mitotic exit to START, START to budding and budding to mitotic exit **(Figure S5)**. The absolute changes in duration between the wild type and the two TAG mutants were comparable for the cell cycle phases and the whole cell cycle and on average were equal to 5 minutes, which corresponds to one imaging interval. Together, these results show that in cell cycles of identical length, START occurs later in *ΔDGA1ΔLRO1* and *ΔTGL3ΔTGL4* than in the wild type. This suggests that START is delayed in cells that cannot synthesise or mobilise TAG, which signifies that intact TAG metabolism is important for the timely occurrence of START.

**Figure 5.**
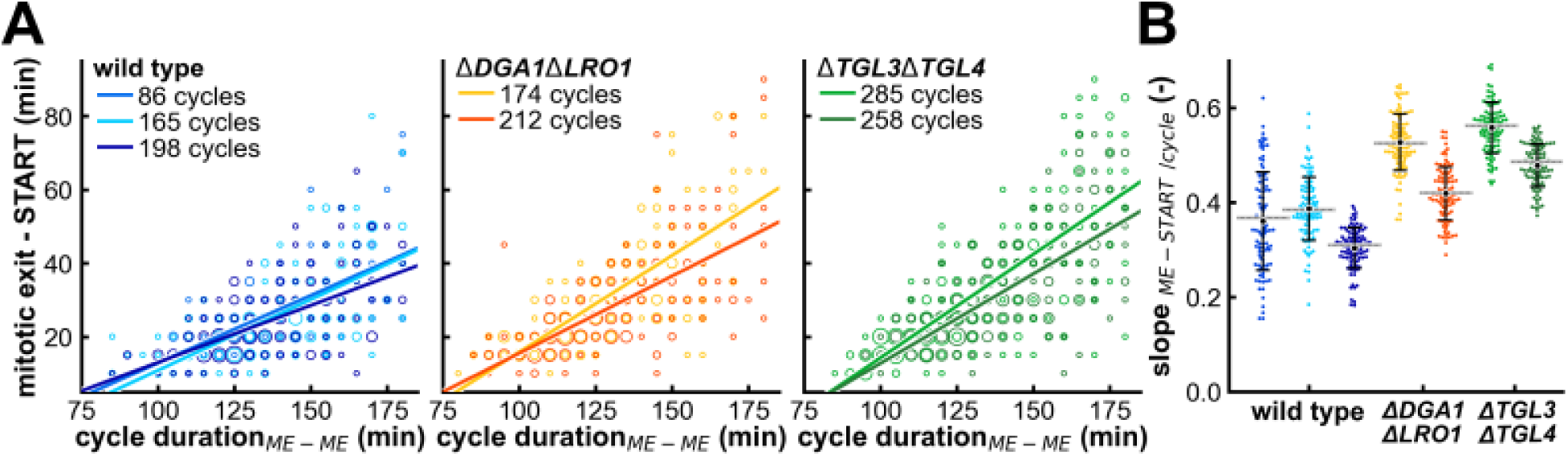
Perturbing LDs through TAG metabolism delays START. **(A)** Interrelation between the duration of mitotic exit to START and duration of the whole cell cycle in the wild type, *ΔDGA1ΔLRO1* and *ΔTGL3ΔTGL4*. Different colours represent replicate experiments and marker size scales with the number of times a combination of cell cycle length and mitotic exit to START duration was observed. Trend lines obtained with linear regression describe the duration of mitotic exit to START duration as a function of cell cycle length; **(B)** Variation in the slope values of the regression lines from A, estimated with bootstrapping. A total of 100 bootstrapping iterations were performed for every experiment. In each iteration, 50% of data points were randomly sampled with replacement and subsequently, linear regression was performed to obtain a slope value. Mean and standard deviation of the slope values obtained with bootstrapping are shown in black. Grey horizontal lines indicate the slope values of the regression lines in A, which were obtained using all data points.

## DISCUSSION

In this work, we established Pln1, Tgl3 and Rrt8 as marker proteins for LDs by showing their colocalisation with LDs stained with BODIPY-TR. Using Pln1-mNeonGreen and mNeonGreen-Tgl3 as LD reporters in time-lapse microscopy experiments, we found that the area-normalised LD number oscillates during the cell cycle, with a minimum around START and a peak halfway through S/G2/M. We perturbed TAG metabolism by deleting the genes that encode the acyl transferases Dga1 and Lro1 or the lipases Tgl3 and Tgl4, respectively preventing the synthesis of TAG or its mobilisation from LDs. Both sets of gene deletions abolished the cell cycle dynamics in the area-normalised LD number. Furthermore, in cell cycles of identical duration, START on average was delayed in the two double deletion backgrounds compared to the wild type, suggesting that the mobilisation of TAGs from LDs is important for START to take place. Thus, our results demonstrate the importance of storage lipid metabolism for cell cycle progression.

The LD cell cycle dynamics that we observed are supported by a number of previous publications, indicating biological processes that may contribute to the dynamic behaviour of LDs. First, cells that re-enter the cell cycle after starvation mobilise neutral lipids from LDs (Kurat et al., 2006; Rajakumari & Daum, 2010). As we have shown here, exponentially growing cells do the same when passing START. Furthermore, both the protein levels of the fatty acid synthesis enzymes Acc1, Fas1 and Fas2 (Blank et al., 2017) and lipid biosynthetic activity (Takhaveev et al., 2023) peak in S/G2/M, which could explain the increase in area-normalised LD number we observed during this part of the cell cycle. Moreover, the lipids that are mobilised from LDs during the second half of S/G2/M could partake in triglyceride cycling, a process in which TAG is partially degraded and then re-synthesised with different fatty acid chains, to metabolically alter stored neutral lipids and change their exact molecular identity (Wunderling et al., 2023). Finally, neutral lipid storage is important right before mitotic exit, as cells that lack all LDs exhibit cytokinetic defects, which are rescued by chemical inhibition of lipid synthesis (P. L. Yang et al., 2016).

However, a recent study that quantified TAG in cells going through the yeast metabolic cycle (YMC) (S. Yang et al., 2025) contradicts the neutral lipid storage dynamics inferred from the LD number in the current work. Yang *et al*. show an increase in TAG levels early in the YMC, followed by a gradual decrease that lasts until the cycle is completed. We believe that the discrepancy between the LD dynamics we report and these TAG dynamics along the YMC could be due to only 40% of these metabolically synchronised cells dividing. Since LDs are dynamic during stationary phase (Hariri et al., 2018; Qiu et al., 2023; Wang et al., 2014) when cells no longer divide, the dividing and non-dividing cells may well have distinct TAG dynamics. In this case, the reported TAG dynamics along the YMC would not match the cell cycle dynamics of TAG, which in turn could explain the differences between the dynamics of TAG (S. Yang et al., 2025) and the cell cycle dynamics of the area-normalised LD number reported in the current work.

Multiple cell cycle regulators could be involved in the orchestration of LD dynamics. The cyclin-dependent kinase Cdc28 activates the lipase Tgl4 (Kurat et al., 2009) and inhibits Pah1 (Choi et al., 2011; Santos-Rosa et al., 2005), which synthesises the TAG precursor diacylglycerol. Pah1 is also regulated by PKA (Su et al., 2018) and PKC (Su et al., 2014). Activity of TORC1 along the cell cycle (Guerra, Vuillemenot, Van Oppen, et al., 2022) coincides with periods of mobilisation of neutral lipids from LDs. Indeed, TORC1 activity has been shown to stimulate the mobilisation of neutral lipids from LDs (Madeira et al., 2014). Thus, the cell cycle regulators Cdc28, PKA, PKC and TORC1 conceivably could participate in the coordination of the cell cycle dynamics of LDs.

We have shown that mobilisation of TAG from LDs contributes to the timely occurrence of START. Previous findings indicate that the mobilised lipids could be used as precursors for sphingolipid synthesis. Supplementation with a sphingolipid precursor rescues the delay in START in *ΔTGL3ΔTGL4* (Chauhan et al., 2015) while inhibition of sphingolipid synthesis inhibits the G1/S transition (Cerbón et al., 2005). Sphingolipids can activate the phosphatase PP2A^Cdc55^ (Nickels & Broach, 1996), which in turn promotes START (McCourt et al., 2013; Moreno-Torres et al., 2015). Thus, sphingolipid synthesis can drive cell cycle progression and lack thereof could explain the delay in START observed in TAG mutant strains.

Overall, we have discovered that the number of LDs oscillates along the cell cycle, that this oscillation depends on TAG metabolism and that the mobilisation of TAG from LDs is important for the timely occurrence of START. Our findings highlight the importance of storage lipid metabolism in cell cycle progression and emphasise that LD storage should be considered in research centred on commitment to the cell cycle. Further research is still needed to elucidate the regulatory processes underlying the observed dynamics of LDs.

## MATERIALS AND METHODS

### Strains

The yeast strains used in this study **(Table S1)** were constructed from the wild-type strain YSBN6 (S288C background) (Canelas et al., 2010) or from YSBN6 *WHI5::mCherry-BLE* (Litsios et al., 2019), using a CRISPR-Cas9 based approach (Novarina et al., 2022) with primers listed in **Table S2**. To tag proteins with the fluorescent protein mNeonGreen, we used a codon-optimised sequence for expression in *S. cerevisiae* (Guerra, Vuillemenot, Rae, et al., 2022). Gene deletions were verified by PCR. Introduction of a fluorescent protein on target proteins was verified by PCR and sequencing.

### Culturing

Cells were grown in 10 mL of Verduyn minimal medium (Verduyn et al., 1992) buffered at pH 5.0 with 10 mM potassium phthalate and with 2% glucose as a carbon source in 100 mL flasks at 30°C, under constant rotation at 300 rpm. To obtain snapshots of live cells, 2-4 µL of cells were taken from an exponentially growing culture and placed on a microscopy slide under a 1% agarose pad soaked with the culture medium. To prepare cells for microfluidic experiments, cultures were maintained in the exponential phase for at least 12 h prior to the experiment through repeated dilution. On the day of the experiment, cultures were diluted to an OD_600_ of 0.05 and after one additional doubling, cells were loaded into the microfluidic chip. Microfluidics experiments were performed as described (Huberts et al., 2013). The flow rate during microfluidics experiments was 4 µL/min.

### Formaldehyde fixation and BODIPY-TR staining

Cells were grown for 24 h to reach stationary phase. The equivalent of 1 mL of a culture with an OD_600_ of 5 was harvested by centrifugation (3 min, 10 000 g). Next, cells were fixed with formaldehyde as described (Madeira et al., 2015). Briefly, cells were resuspended in 1 mL of 3.7% (w/v) paraformaldehyde with 0.1 M sorbitol as an osmolyte and incubated at room temperature for 15 min. Afterwards, cells were washed once with 1 mL of phosphate buffered saline (PBS) and finally resuspended in 1 mL of PBS.

To stain formaldehyde-fixed cells with BODIPY-TR methyl ester (Lumiprobe GmbH) 1 µL of 5 mM BODIPY-TR methyl ester dissolved in DMSO was added to 100 µL of fixed cell suspension and incubated at room temperature for 5 min. Cells were then pelleted (1.5 min, 12 500 g), washed once with 100 µL of PBS and finally resuspended in 100 µL of PBS. Cells were imaged immediately after staining.

### Microscopy

All images were acquired on a Nikon Eclipse Ti-E inverted wide-field fluorescence microscope equipped with the Nikon Perfect Focus System (PFS) and an Andor-DU-897 EX camera. For imaging, a 100x S Fluor Oil objective (NA 1.4, Nikon) was used and the readout mode was set to 1 MHz without gain amplification. For bright-field images, excitation was done with a halogen lamp fitted with a 420 nm beam-splitter to filter out short wavelengths. A Lumencor AURA excitation system was used for fluorescence excitation. Green fluorescent proteins were excited at 485 nm with an imaging set-up consisting of a 470/40 nm band-pass filter, a 495 nm beam splitter and a 525/50 nm emission filter. Red fluorophores were imaged using excitation at 565 nm, a 560/40 nm band-pass filter, a 585 nm beam splitter and a 630 nm/75 nm emission filter. Intensity of the light source and excitation time for each fluorophore and each experiment are detailed in **Table S3**. For live-cell imaging, the microscope setup was kept at 30 °C using an incubator box (Life Imaging Services).

### Optimising image alignment

To correct the small-scale but systematic misalignment between images recorded in the GFP channel (mNeonGreen-tagged proteins) and the RFP channel (BODIPY-TR), we used the bilinear transformation approach from the Python module pyStackReg (Thévenaz et al., 1998). To obtain the transformation matrix, we created composite images of binarised snapshot images of cells that express Pln1-mNeonGreen and that were also stained with the red fluorophore BODIPY-TR. Binarisation of the snapshot images ensured that only outlines of the cells, but no cytoplasmic structures, were visible. Combination of multiple snapshot images yielded a composite image showing the outline of cells dispersed over the whole field of view. We aligned the composite of RFP snapshots to the composite of the corresponding GFP snapshots, thereby obtaining a transformation matrix which we used to optimally align images recorded in the RFP channel to images recorded in the GFP channel.

### Cell segmentation and cell cycle alignment

Fluorescence images were background corrected using the rolling ball algorithm implemented in ImageJ with a radius of 50 pixels. Cell masks were obtained from bright-field images with the ImageJ plugin BudJ (Ferrezuelo et al., 2012). Segmentation was inspected visually to verify that the cell masks matched the cells in the bright-field images. Finally, cell cycles were excluded from further analysis if their total duration, from one mitotic exit to the next, exceeded 180 min. For all cells, we manually tracked mitotic exit, budding and START. Budding was detected in the bright-field images, START was detected from Whi5-mCherry leaving the nucleus and mitotic exit from Whi5-mCherry entering the nucleus.

For the cell cycles that passed the selection criteria (*i.e.* proper segmentation and cell cycle duration), we transformed the data onto a common cell cycle progression coordinate ranging from 0 to 1 to allow comparison between cycles of different durations. Consecutive mitotic exits were defined as 0 and 1. We also determined the average position of START and budding on this common time coordinate. To do so, we calculated the fraction of the cell cycle that had passed at the moment that each event occurred. Specifically, to determine the timing of START, we divided the time between the first mitotic exit and START by the duration of the whole cell cycle. We repeated this procedure for budding. Ultimately, combining data from all cell cycles, we determined the average timing of START and budding on the normalised cell cycle progression coordinate.

Subsequently, we determined the cell cycle progression coordinate value for every data point recorded in the time-lapse microscopy experiments. To do this, we divided each cycle into three phases: mitotic exit to START, START to budding and budding to mitotic exit. For each of these phases, we placed its first and last data point, which coincide with a cell cycle event, at the normalised time values for these events. Then, we evenly dispersed all interstitial data points over the time interval bounded by the two cell cycle events. Performing this procedure for every cell cycle, we determined a time value on the common cell cycle progression coordinate for every recorded data point.

To infer the population-average behaviour of the various measures over the cell cycle, we performed Gaussian process regression of the cell cycle aligned single-cell data. For this, we used Python’s `sklearn.gaussian_process` (Pedregosa et al., 2011) using the radial basis function (RBF) as a prior, with the length scale range [0.01, 0.5] and the white kernel with free noise level. An optimised fit was obtained through maximisation of the log-marginal likelihood in the regression.

### Automated detection of LDs in fluorescence microscopy images

We used the PunctaFinder algorithm (Terpstra et al., 2024) to automatically detect LD puncta and estimate their size, both for LDs stained with BODIPY-TR in the RFP channel and for LDs visualised with fluorescently tagged marker proteins in the GFP channel. We used an overlap parameter value of zero and a punctum diameter of three pixels, which reflects a distance of 480 nm on our microscopy setup and corresponds to the diameter of an average lipid droplet (*i.e.* 400 nm) (Czabany et al., 2008). To obtain suitable threshold values for punctum detection, we created a manually curated data set of 40 cells for each genetic background and performed threshold value optimisation with five bootstrap iterations, sampling 75% of cells with replacement. Each final threshold value was the average of the values obtained in the five bootstrap iterations. The threshold values are provided in **Table S4**.

### Colocalisation analysis

Colocalisation between puncta of LDs stained with BODIPY-TR and puncta of LD marker proteins tagged with mNeonGreen was assessed based on the distance between the midpoints of puncta in the two imaging channels. Puncta are qualified as colocalising if their midpoints are maximally one pixel apart in both the x-and y-direction. Colocalisation is qualified as ambiguous for puncta whose midpoints are two pixels apart in one direction, but maximally one pixel apart in the other. In all other cases, colocalisation is qualified as non-existent.

## SUPPLEMENTARY INFORMATION

**Figure S1.**
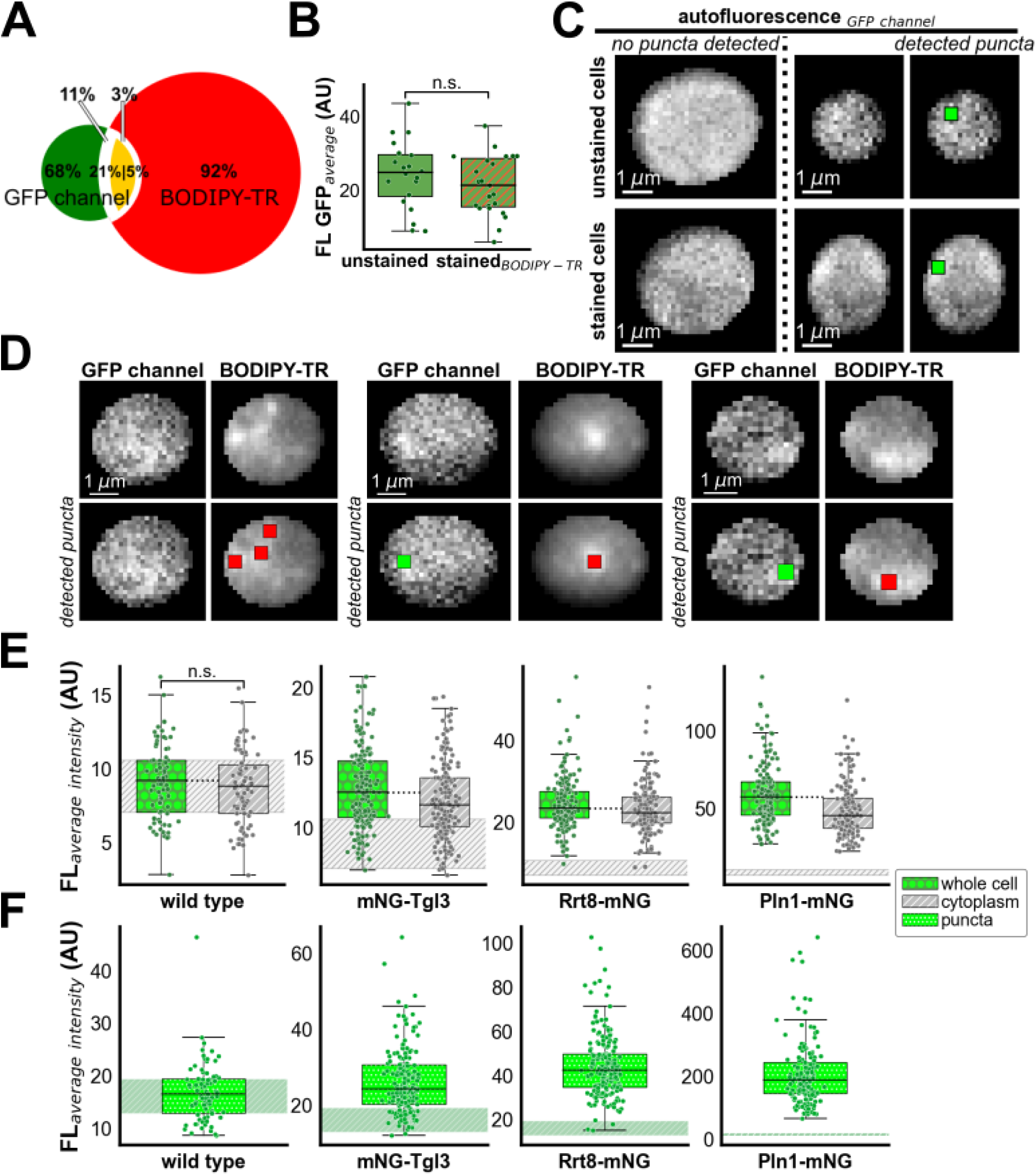
BODIPY-TR fluorescence is not detected in the GFP channel and puncta of GFP autofluorescence are not detected in cells expressing an LD reporter tagged with mNeonGreen. **(A)** Colocalisation between LDs stained with BODIPY-TR and puncta detected in the GFP channel in wild-type cells. The left circle of the Venn diagram represents all LDs identified with BODIPY-TR in the RFP channel; the right circle represents all puncta identified in the GFP channel. The orange overlapping region are puncta that colocalise between the LDs stained with BODIPY-TR and the puncta in the GFP channel. The red region indicates LDs that do not colocalise with a punctum in the GFP channel, while the green region indicates puncta in the GFP channel that do not colocalise with an LD stained with BODIPY-TR. The white regions represent puncta with ambiguous colocalisation: colocalisation between puncta in the two fluorescence channels is only probable, not certain, despite their midpoints being close together. 274 cells were analysed, 427 BODIPY-TR puncta and 89 puncta in the GFP channel were identified; **(B)** Comparison of the average fluorescence intensity measured in the GFP channel for cells stained with BODIPY-TR and unstained cells. Since the cells stained with BODIPY-TR are not brighter than the unstained cells, we can conclude that BODIPY-TR fluorescence is not detected in the GFP channel; **(C)** Fluorescence microscopy images of wild-type cells, recorded in the GFP channel. The top row shows unstained cells, while the bottom row shows cells stained with the red fluorophore BODIPY-TR. There are no clearly visible differences between stained and unstained cells, demonstrating that BODIPY-TR staining does not influence images recorded in the GFP channel. Furthermore, puncta could be detected in images of both stained and unstained cells, as shown in the third column of images. This finding indicates that brighter spots in the autofluorescence occur naturally and can result in the appearance of puncta; **(D)** Fluorescence microscopy images and detected puncta of three wild-type cells stained with BODIPY-TR. The puncta identified in the GFP channel images are only slightly brighter than the cytoplasm and, in the visualised cells as well as the majority of other cells, do not colocalise with BODIPY-TR puncta, ruling out that these GFP puncta are due to detection of BODIPY-TR signal in the GFP channel; **(E-F)** Average fluorescence intensity of the cytoplasm of cells with at least one punctum, the same whole cells, *i.e.* cytoplasm and puncta combined, and the detected puncta. Shaded areas in E and F indicate the box (first quartile to third quartile) of whole wild-type cells or puncta detected in wild-type cells, respectively. The punctum detection threshold values to detect GFP puncta in wild-type cells are more stringent than those applied to cells expressing mNG-Tgl3 or Rrt8-mNG **(Table S4)**. Still, the average fluorescence intensity of the cytoplasm is significantly lower than that of whole cells in the three reporter strains, but not in the wild-type control (one-sided Mann-Whitney U-test, p<0.01). This finding indicates that the GFP autofluorescence puncta detected in the wild type are similar to the cytoplasm with regards to fluorescence intensity, while in the three LD reporter strains, the bright puncta cause the average fluorescence intensity of whole cells to be higher than that of the cytoplasmic region alone. Moreover, the average fluorescence intensity of the GFP autofluorescence puncta detected in the wild type is notably lower than that of puncta detected in any of the three LD reporter strains. Together, these results indicate that it is improbable that GFP autofluorescence puncta are detected in the three LD reporter strains.

**Figure S2.**
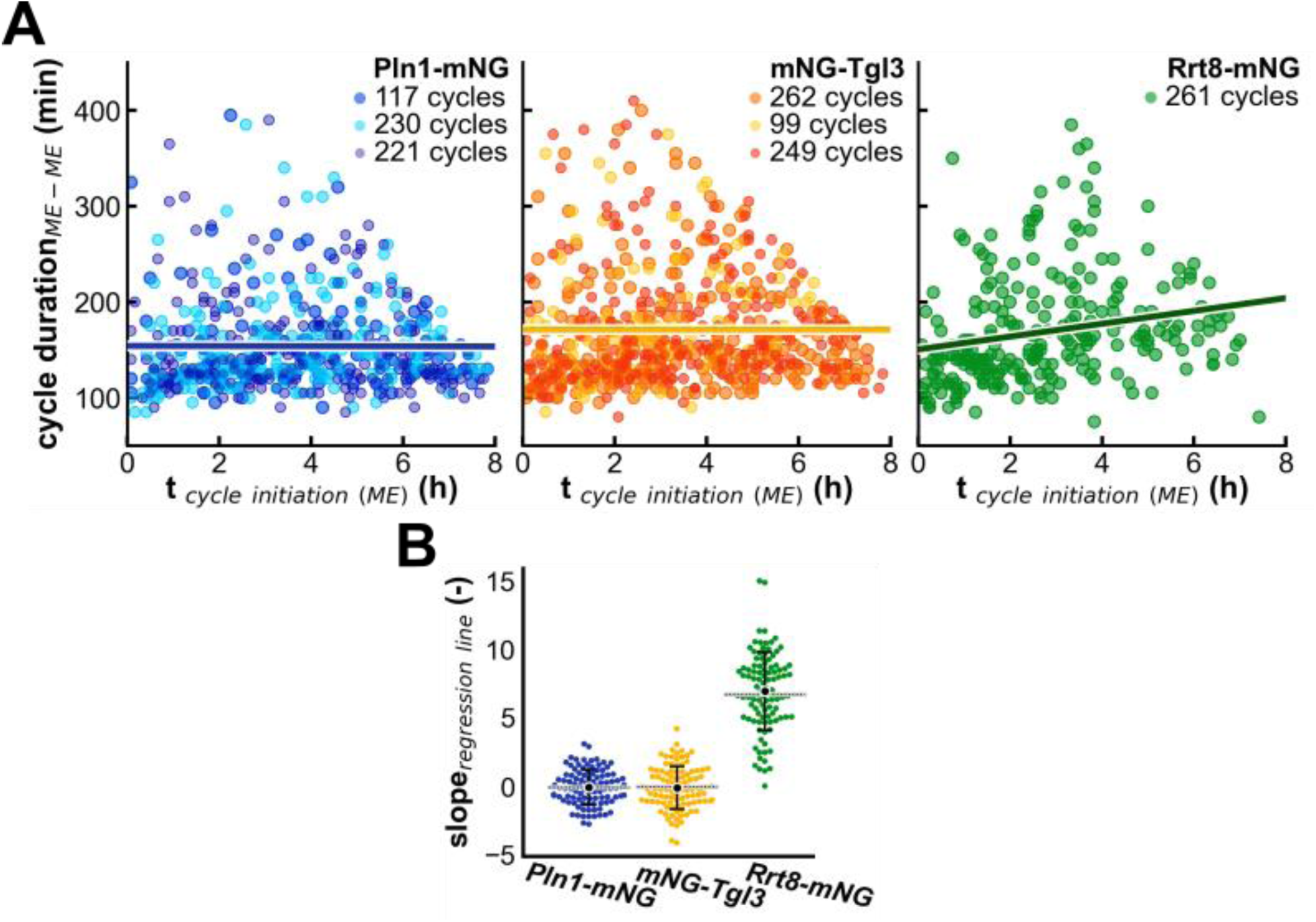
Cell cycle length increases with experiment duration in time-lapse imaging of Rrt8-mNG but not Pln1-mNG and mNG-Tgl3. **(A)** Cell cycle length, from one instance of mitotic exit (ME) to the next was plotted against the starting time of the cycle within the time-lapse experiment for cells expressing Pln1-mNG, mNG-Tgl3 or Rrt8-mNG as a reporter protein for LDs. Replicate experiments are shown with distinct colours. Trend lines describing cell cycle length as a function of cycle initiation time were obtained with linear regression, which was performed on pooled data of replicate experiments; **(B)** Bootstrapping was performed to estimate the variation in the slope of the regression lines from A. For each bootstrap iteration, 50% of the data was randomly sampled with replacement and linear regression was performed to obtain a slope value; a total of 100 iterations were performed for each genetic background. Grey horizontal lines indicate the slope values of the regression lines in A, which were obtained using all data points. Mean and standard deviation of the slopes obtained with bootstrapping are indicated in black. Interestingly, regression lines fitted to data from cells expressing Rrt8-mNG have positive slope of values, while regression lines fitted to data from cells expressing Pln1-mNG or mNG-Tgl3 have an average slope value of approximately 0. Thus, in time-lapse imaging of cells expressing Rrt8-mNG cell cycle length increases the longer the experiment has lasted. This reveals a potential phototoxic effect, caused by the cumulative effects of repeated exposure to the lasers used for fluorophore excitation.

**Figure S3.**
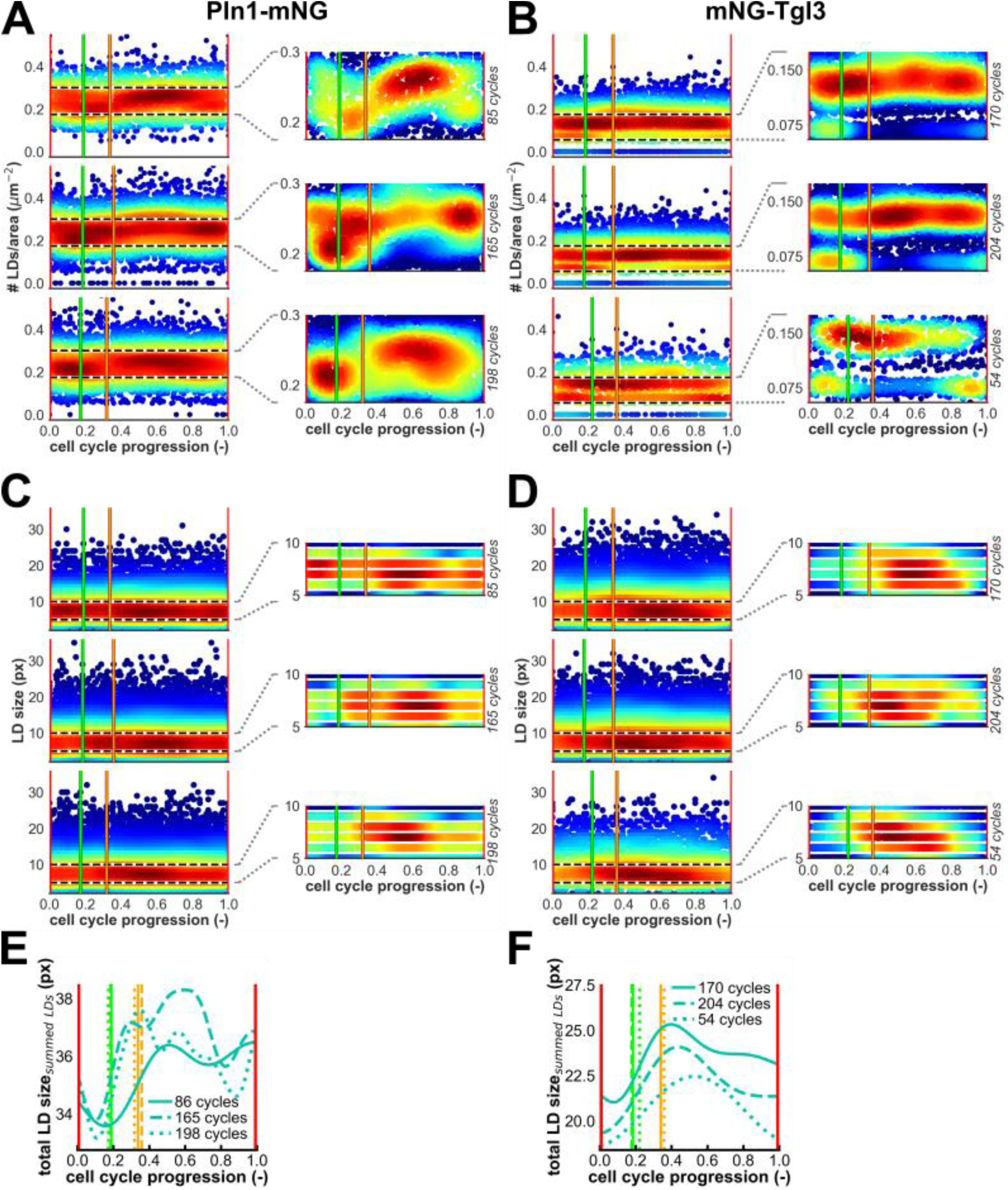
LD number is dynamic along the cell cycle while LD size is constant. LDs were identified in time-lapse microscopy images of cells expressing either Pln1-mNG (A, C, E) or mNG-Tgl3 (B, D, F) as an LD marker protein. For both reporter strains, three replicate experiments were performed. Cell cycle trajectories were aligned from one occurrence of mitotic exit (ME) to the next (red vertical lines at cell cycle progression values 0 and 1) and were also aligned for START (bright green vertical line) and budding (orange vertical line). **(A-D)** Density estimations, showing densely populated data points in red and sparsely populated data points in blue, obtained with Gaussian kernel estimation, show the cell cycle dynamics of (A, B) the number of LDs normalised to the cell cross-area and (C, D) LD size. The number of cell cycles assessed in each replicate is indicated on the right side of each plot. The plots in each second column zoom in on the plots with all data and show their most densely populated region, located between the dashed horizontal lines. Here, the colour map has been rescaled to assess the density in more detail. Notably, the density plots show the same cell cycle dynamics of the LD number normalised to the cell cross-area predicted with Gaussian process regression (Figure 2A) when Pln1-mNG is used as an LD reporter protein, but not with mNG-Tgl3. Still, with mNG-Tgl3 as an LD marker, relatively dense subpopulations with <0.1 LDs/µm^2^ are visible early in the cell cycle in all three replicates, reflecting the trough around START in the cell cycle dynamics of the number of LDs per cell cross-area predicted with Gaussian process regression (Figure 2B). Also, with mNG-Tgl3 as an LD reporter protein, distinct subpopulations of cells with one or two puncta are visible, at #LD/area values of approximately 0.07 LDs/µm^-2^ and 0.14 LDs/µm^-2^, respectively. These subpopulations occur since the area-normalised LD number is obtained by dividing the discrete number of LDs by the continuous cell cross-area. Evidently, the range in cell cross-area values is narrow, causing the resulting area-normalised LD number to still appear discrete in the density plots; **(E-F)** Gaussian process regression outputs showing the cell cycle dynamics of the total area of detected LDs, *i.e.* summed sizes of all LDs detected in a cell, without normalisation to the cell cross-area. Since these outputs resemble the cell cycle dynamics of the area-normalised LD number, its oscillation does not result from the normalisation to the cell cross-area. Moreover, the strong resemblance between the cell cycle dynamics of the summed LD size and the dynamics of the area-normalised LD number further confirms that LD number, but not LD size, is dynamic along the cell cycle.

**Figure S4.**
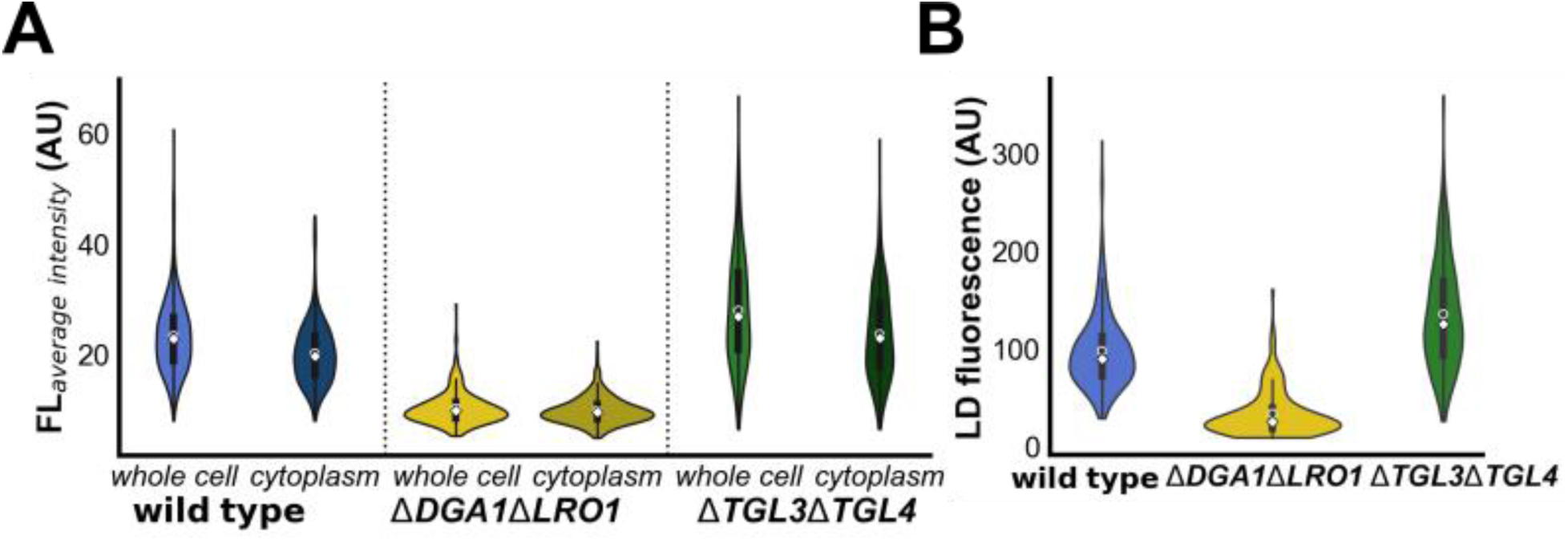
Genetic perturbation of TAG metabolism affects Pln1-mNG expression levels. TAG metabolism was perturbed by gene deletion of either *DGA1* and *LRO1*, which encode the main TAG synthases, or *TGL3* and *TGL4*, which encode the lipases responsible for TAG mobilisation from LDs. Average Pln1-mNeonGreen fluorescence intensity was determined in snapshot images of cells from an exponential culture for **(A)** the whole cell and the cytoplasmic region as well as **(B)** the LDs in the wild type, *ΔDGA1ΔLRO1* and *ΔTGL3ΔTGL4*. White diamonds indicate the median and open circles indicate the population average. Both genetic perturbations led to significant changes in measured Pln1-mNeonGreen fluorescence compared to the wild type (two-sided Mann-Whitney U-test, p < 0.001) for all three regions of interest. Notably, the change in Pln1-mNG fluorescence compared to the wild type is much larger for *ΔDGA1ΔLRO1* than for *ΔTGL3ΔTGL4*, as quantified using Cohen’s d to assess the effect size. For *ΔDGA1ΔLRO1*, the effect size for the changing fluorescence intensity between deletion strain and wild type was 2.54, 2.40 and 1.86 for the whole cell, the cytoplasm and the puncta, respectively. In contrast, for *ΔTGL3ΔTGL4*, the effect size for these three regions of interest was equal to 0.54, 0.49 and 0.79.

**Figure S5.**
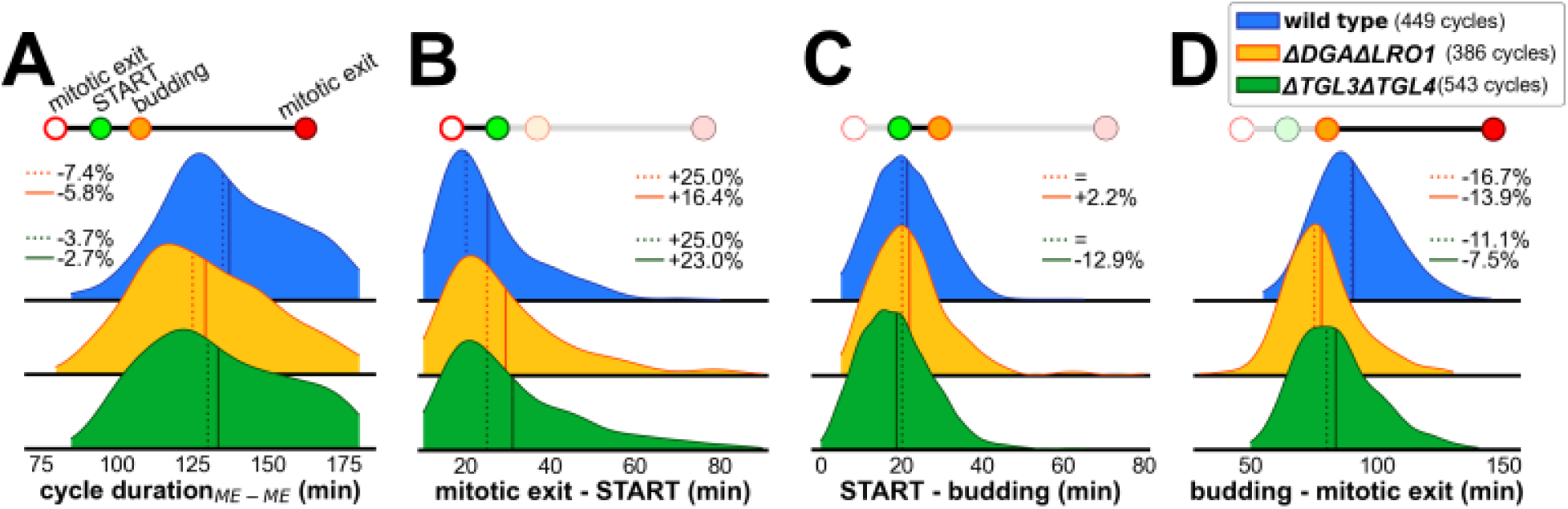
Duration of the cell cycle and its subphases are altered slightly in ΔDGA1ΔLRO1 and ΔTGL3ΔTGL4 compared to the wild type. Probability density functions for duration of **(A)** the whole cell cycle and the cell cycle phases **(B)** mitotic exit to START, **(C)** START to budding and **(D)** budding to mitotic exit in the wild type, *ΔDGA1ΔLRO1* and *ΔTGL3ΔTGL4*. To obtain these distributions, cell cycles recorded in three replicate experiments with the wild type and two replicate experiments each with *ΔDGA1ΔLRO1* and *ΔTGL3ΔTGL4* were pooled. Solid and dotted vertical lines denote the mean and median value of each distribution, respectively. Percentages denote the change in median and mean in each deletion strain compared to the wild type. Above each plot, a schematic representation of the cell cycle indicates the cell cycle phase(s) studied. Notably, only cell cycles with a total duration of at most 180 min were included in the analysis.

**Table S1.** Yeast strains used in the current study.

| Strain | Source |
| --- | --- |
| YSBN6 (S288C-derived strain, MATa FY3 HO::HphMX4) | Canelas *et al.*, 2010 |
| YSBN6 *WHI5::mCherry-BLE* | Litsios *et al.*, 2021 |
| YSBN6 *PLN1::mNeonGreen* | This study |
| YSBN6, *TGL3::mNeonGreen-Tgl3* | This study |
| YSBN6 *RRT8::mNeonGreen* | This study |
| YSBN6 *WHI5::mCherry-BLE PLN1::mNeonGreen* | This study |
| YSBN6 *WHI5::mCherry-BLE TGL3::mNeonGreen-Tgl3* | This study |
| YSBN6 *WHI5::mCherry-BLE RRT8::mNeonGreen* | This study |
| YSBN6 *WHI5::mCherry-BLE PLN1::mNeonGreen ΔDga1 ΔLro1* | This study |
| YSBN6 *WHI5::mCherry-BLE PLN1::mNeonGreen ΔTgl3 ΔTgl4* | This study |

**Table S2.**
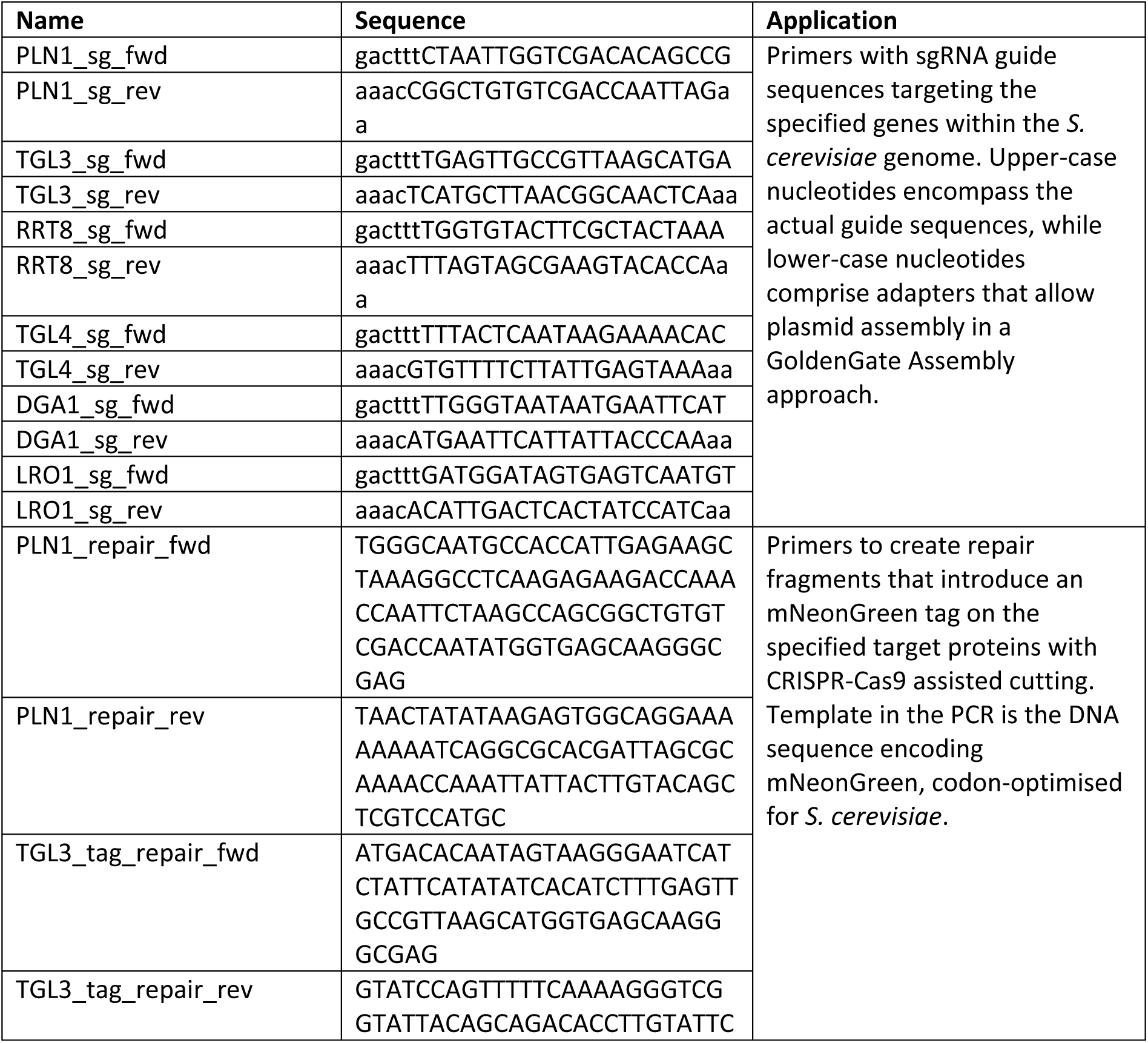
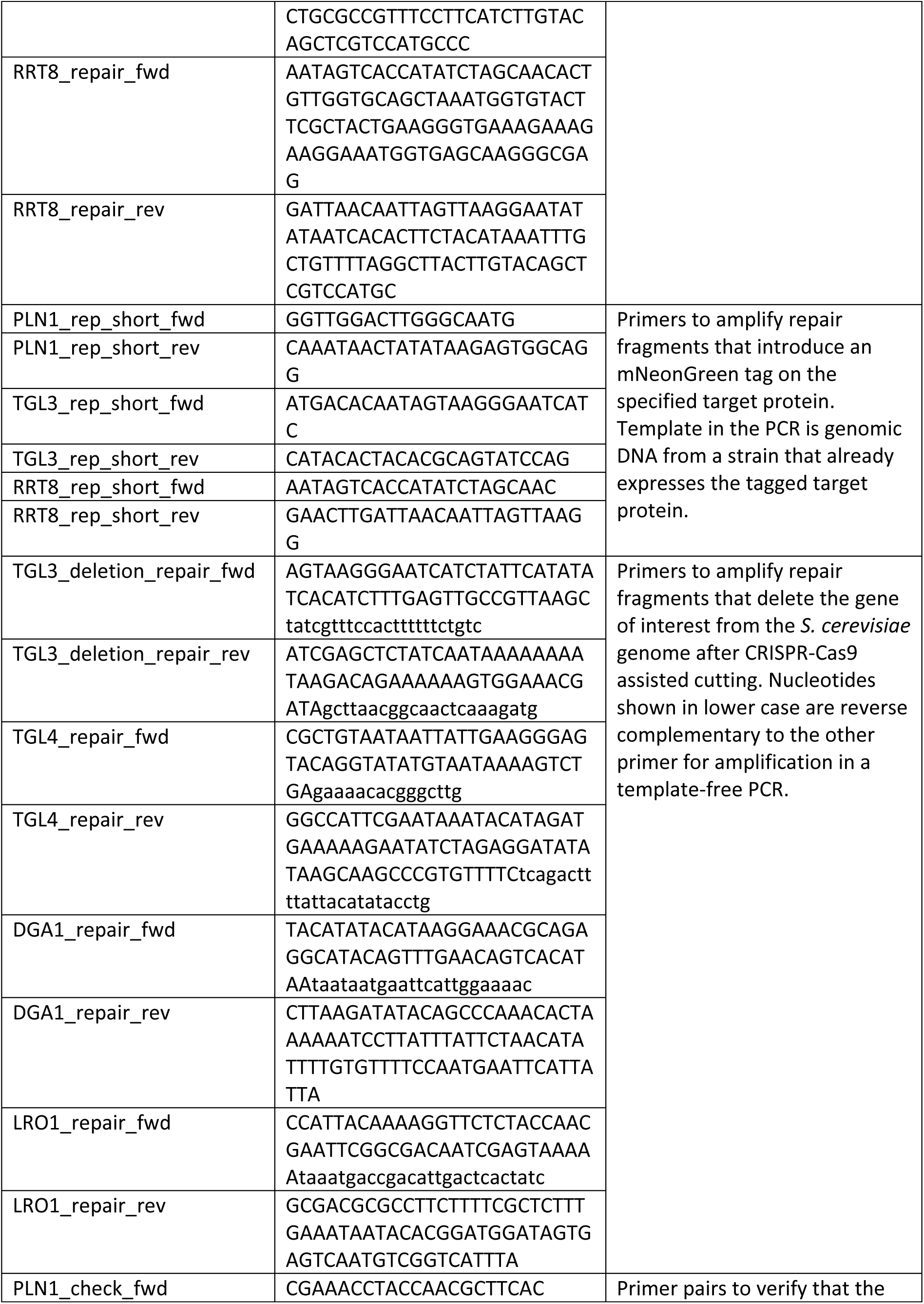
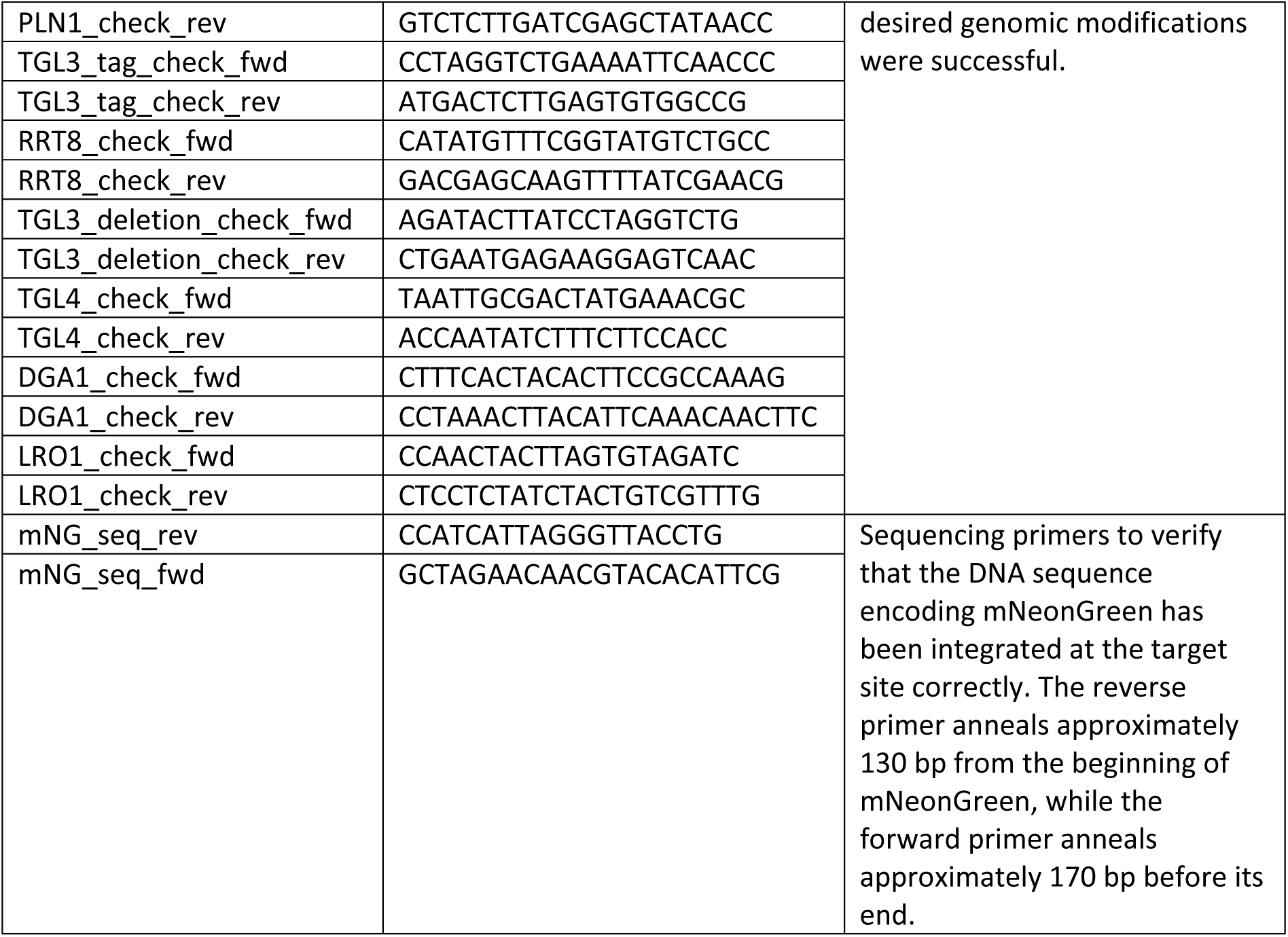
Primers used in the current study.

**Table S3.** Imaging settings used for microscopy experiments. For each fluorophore and protein of interest, the imaging channel, light intensity and excitation time applied are detailed below.

| Target | Imaging channel | Light intensity | Excitation time |
| --- | --- | --- | --- |
| BODIPY-TR (fixed cells) | RFP | 1% | 5 ms |
| Pln1-mNG (fixed cells) | GFP | 7% | 300 ms |
| mNG-Tgl3 (fixed cells) | GFP | 7% | 300 ms |
| Rrt8-mNG (fixed cells) | GFP | 7% | 300 ms |
| autofluorescence (fixed cells) | GFP | 7% | 300 ms |
| Whi5-mCherry (live cells; time-lapse) | RFP | 10% | 300 ms |
| Pln1-mNG (live cells; time-lapse) | GFP | 3% | 200 ms |
| mNG-Tgl3 (live cells; time-lapse) | GFP | 3% | 200 ms |

**Table S4.** Threshold values used for automated punctum detection with PunctaFinder. Threshold values for punctum detection in fluorescence microscopy images of the neutral lipid dye BODIPY-TR (RFP channel) and mNeonGreen (mNG) tagged reporter proteins for LDs or a wild-type autofluorescence control (GFP channel). Because of slight differences in BODIPY-TR staining between the four samples, separate thresholds were determined for punctum detection in the RFP channel as well. In all cases, the punctum diameter was set to three pixels and the overlap parameter value to zero. Thresholds were determined based on manually validated datasets of 42 cells expressing Pln1-mNG, 41 cells expressing mNG-Tgl3, 43 cells expressing Rrt8-mNG and 45 wild-type cells.

| **Genetic background** | **Fluorescence channel** | **T<sub>ratio, local</sub>** | **T<sub>ratio, global</sub>** | **T<sub>cv</sub>** |
| --- | --- | --- | --- | --- |
| Pln1-mNG | GFP | 1.408 | 1.406 | 0.358 |
|  | RFP | 1.252 | 1.338 | 0.262 |
| mNG-Tgl3 | GFP | 1.342 | 1.508 | 0.272 |
|  | RFP | 1.264 | 1.364 | 0.252 |
| Rrt8-mNG | GFP | 1.348 | 1.353 | 0.242 |
|  | RFP | 1.254 | 1.398 | 0.206 |
| wild type | GFP | 1.380 | 1.648 | 0.266 |
|  | RFP | 1.222 | 1.382 | 0.194 |

